# Computational Analysis of Zn^2+^ Mediated Non-Competitive Inhibition to Caspase-3

**DOI:** 10.1101/2024.09.30.615770

**Authors:** Xinyu Erya Tian, Min Zang, Xiaoyu Wang, Hao Dong

## Abstract

Caspase-3, a critical mediator of apoptosis, is implicated in neurodegenerative diseases and cancer, making it an attractive therapeutic target. Here, we elucidated the molecular mechanisms of caspase-3 inhibition through computational methods. We first investigated the binding of the canonical substrate DEVD and novel peptidomimetic inhibitors, characterizing their competitive inhibition mechanisms. Using AlphaFold3 and molecular dynamics simulations, we then identify Zn^2+^-binding sites and characterize conformational changes induced by Zn^2+^ chelation, revealing a mechanism for non-competitive inhibition. Our results provide atomic-level insights into both competitive and Zn^2+^-mediated non-competitive inhibition of caspase-3, enhancing our understanding of apoptotic pathways. These findings could help guide the development of targeted therapies for apoptosis-related disorders, encompassing both competitive inhibitors and non-competitive modulators.

## 1. Introduction

Caspases, pivotal enzymes in apoptotic pathways, play a critical role in cellular homeostasis. Dysregulation of these proteases is intricately linked to the pathogenesis of various disorders, particularly neurodegenerative conditions such as Alzheimer’s and Parkinson’s diseases, as well as cancer progression.^1-3^ Aberrant expression or activity of caspases often proves crucial in pathological processes. Elucidation of these enzymes has yielded significant insights into disease mechanisms and potential therapeutic interventions, especially in oncology, where modulation of caspase activity may facilitate targeted approaches for apoptosis-related disorders.^4, 5^

All caspases rely on two essential catalytic residues - cysteine (Cys) and histidine (His)— which are indispensable for substrate binding and catalytic cleavage.^6, 7^ These highly conserved residues and structural features enable caspases to execute precise regulatory functions during apoptosis. Numerous caspase types have been identified to date.^8^ Despite variations in their activity and substrate specificity, their overall reaction mechanisms exhibit remarkable similarity. All caspases recognize specific tetrapeptide sequences and cleave at the peptide bond following aspartate (Asp) residues, underscoring their indispensable role in modulating apoptotic signaling cascades. Comparative analysis of the three-dimensional (3D) structures of caspase-3, -7, and -8 has revealed their high homology and structural similarity.^9-11^ These enzymes form heterodimers with each monomer comprising large and small subunits. The core structure consists of six *β*-strands surrounded by five *α*-helices, forming a stable spatial configuration.^12, 13^

A notable example is caspase-3, a key protease frequently activated in early apoptotic processes. In healthy cells, it primarily exists as an inactive precursor and is widely recognized as a major biomarker in apoptosis-related research.^14^ Caspase-3 functions as a Zn^2+^-mediated cysteine protease, inherently inhibited by Zn^2+^ ions. Compounds such as PAC-1, which chelate Zn^2+^, can remove these inhibitory divalent metals, thereby fully activating the enzyme. The active form of caspase-3 specifically targets the tetrapeptide motif Asp-Glu-Val-Asp (DEVD) and hydrolyses the peptide bond at the C-terminus of the aspartate residue. To date, the inhibition mechanism of caspase-3 by Zn^2+^ ions or other inhibitors has been inferred primarily from indirect evidence derived from structural information of enzyme/inhibitor complexes.^15, 16^ To better elucidate the molecular mechanisms of caspase-3 inhibition by various inhibitors and the resultant conformational changes, molecular simulation studies are imperative.

Building upon these foundational insights, our study employs a multifaceted approach to elucidate the molecular mechanisms underlying caspase-3 inhibition. We first investigate the inhibition of caspase-3 by its canonical substrate, DEVD (Asp-Glu-Val-Asp), and several distinct peptidomimetic inhibitors, calculating binding energies to elucidate their competitive inhibitory mechanisms. Furthermore, we utilize AlphaFold3, ^17^ a state-of-the-art artificial intelligence-driven protein structure prediction tool, to identify potential Zn^2+^-binding sites on caspase-3. Subsequent molecular dynamics simulations explore the conformational changes and free energy landscape alterations induced by Zn^2+^ chelation, providing atomistic insights into the Zn^2+^-mediated inhibition mechanism. This comprehensive approach aims to unveil the diverse strategies of caspase-3 inhibition, encompassing both competitive and non-competitive mechanisms. Our findings could potentially inform the development of novel therapeutic interventions targeting this crucial apoptotic enzyme, with implications for both competitive inhibitors and non-competitive modulators that may exploit the Zn^2+^-mediated non-competitive inhibition mechanism.

## 2. Results and Discussion

### 2.1 Competitive inhibitor inhibition mechanism of caspase-3

To elucidate the inhibitory mechanisms of caspase-3 inhibitors, we first investigated competitive inhibitors. We performed molecular docking simulations of three previously identified competitive inhibitors to the DEVD binding pocket, followed by extensive molecular dynamics simulations. Analysis of the backbone root-mean-square deviation (RMSD) revealed stable binding of all three inhibitors to the DEVD pocket throughout the simulation trajectories, indicating robust and persistent interactions.

To quantify the binding affinity of these inhibitors, we employed the Molecular Mechanics Generalized Born Surface Area (MMGBSA) method to calculate binding free energies. Our results demonstrated that these inhibitors exhibit binding affinities comparable to or stronger than the canonical DEVD substrate. Specifically, the calculated binding free energies were -55.94, -46.93, and -45.09 kcal/mol for inhibitors Z-DEVD-FMK, Z-VAD(Ome)-FMK, and Q-VD-Oph, respectively, compared to -44.06 kcal/mol for DEVD (**Figure 1**). These computational findings corroborate the competitive inhibition mechanism of these compounds, as they effectively occupy the substrate-binding site with high affinity. The strong binding energies suggest that these inhibitors can successfully compete with the natural substrate for the active site, thereby inhibiting caspase-3 activity. Furthermore, detailed analysis of the binding modes revealed key protein-ligand interactions that contribute to the inhibitors’ potency. We observed critical hydrogen bonding patterns and hydrophobic contacts that mimic those of the DEVD substrate, explaining their competitive nature. These structural insights provide valuable information for future optimization of caspase-3 inhibitors, potentially guiding the design of more potent and selective compounds.

**Figure 1.**
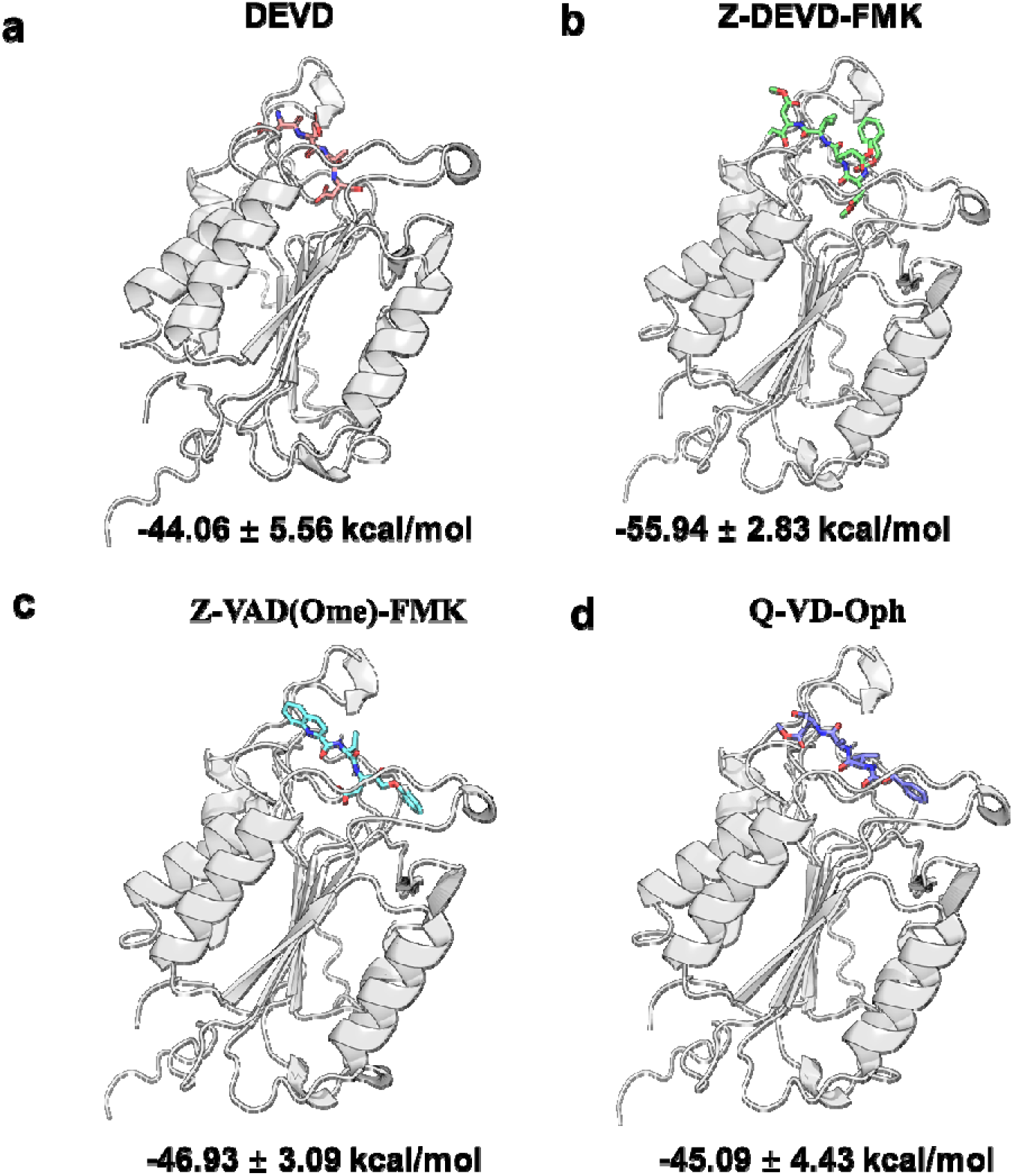
Molecular model diagram, including eutectic structure and molecular docking structure. (a) Cocrystal structure of DEVD and caspase-3. (c-d) Molecular docking structure of inhibitor and caspase-3.

### 2.2 Non-competitive inhibitory sites for Zn^2+^ ion binding of caspase-3

In addition to competitive inhibition, caspase-3 is also subject to non-competitive inhibition by Zn^2+^ ions. Previous investigations have demonstrated that Zn^2+^ inhibition of caspase-3 is predominantly non-competitive in nature. Substrate binding sites remain unaffected by Zn^2+^ ions, with inhibition most pronounced at sub-micromolar concentrations. This observation precludes Zn^2+^ binding to the catalytic dyad. Notably, a crystal structure of the Zn^2+^-caspase-3 complex remains elusive. While the catalytic dyad of the caspase family is postulated to be a common inhibitory Zn^2+^ binding site for all caspases, weakly bound Zn^2+^ may not effectively compete with substrates at the active site. Computational chemistry has corroborated the existence of multiple Zn^2+^ binding sites within caspase-3.

To further elucidate the Zn^2+^-binding mechanism, we employed AlphaFold3, a state-of-the-art artificial intelligence-driven protein structure prediction tool, to predict potential Zn^2+^ ion binding sites on caspase-3. Our analysis revealed a putative binding site coordinated by His121 and Cys163, with additional coordination likely provided by water molecules (**Figure 2**). These findings are consistent with previous computational studies on caspase-3 and show striking similarity to the Zn^2+^-binding site observed in the homologous protein caspase-6.^15^

**Figure 2.**
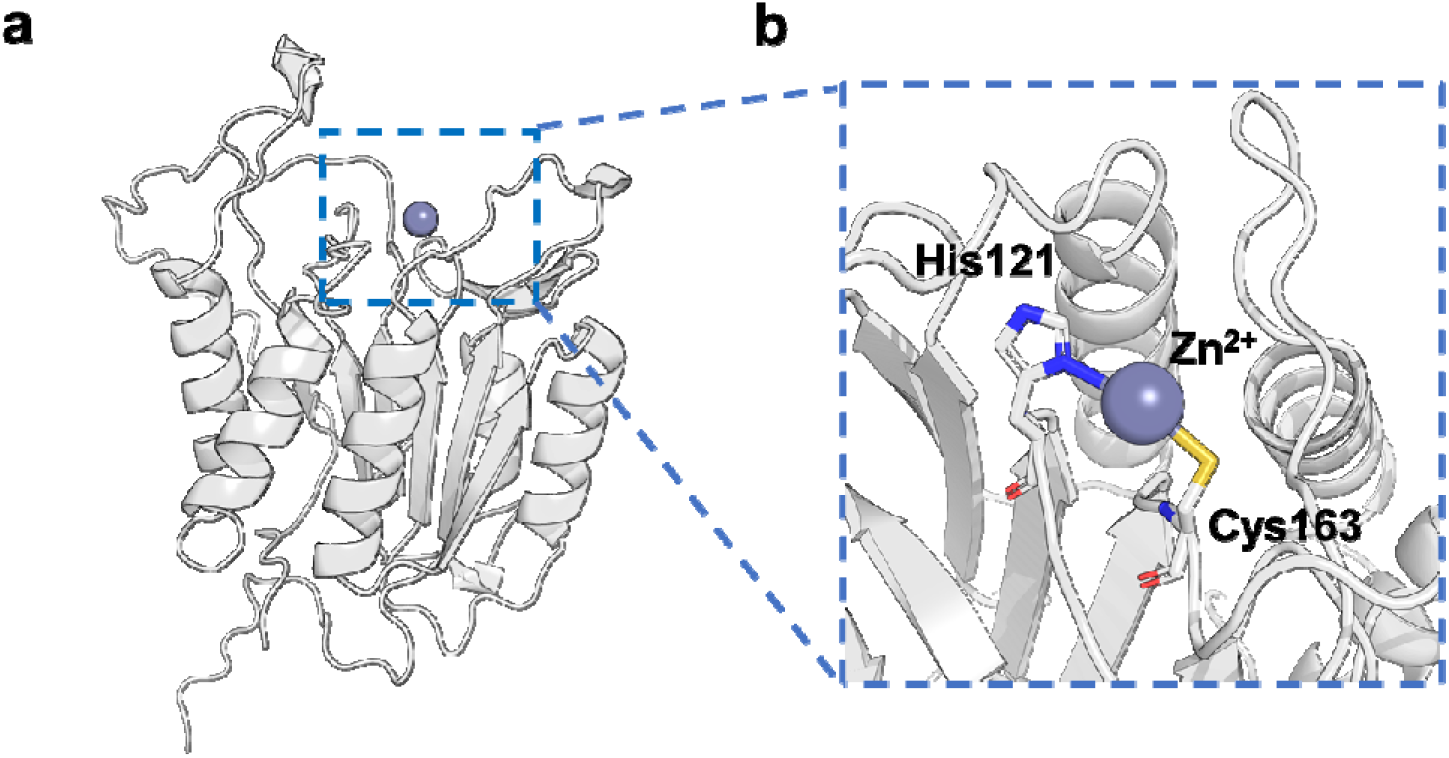
(a) Zn^2+^ ion binding sites predicted by Alphafold3. (b) Zn^2+^ ion chelation with His121 and Cys163 coordination in caspase-3 active pocket.

### 2.3 Interplay between peptide competitive inhibition and Zn^2+^-mediated non-competitive inhibition to Caspase-3

Building upon our prediction of Zn^2+^ binding sites, we further explored the interplay between peptide inhibitors and Zn^2+^-induced effects. We constructed free energy landscape distributions for caspase-3 in both Zn^2+^-bound and Zn^2+^-free states (**Figure 3**). These landscapes, derived from extensive molecular dynamics simulations, provided a comprehensive view of the protein’s conformational space. The distributions unveiled distinct conformational preferences between the two states, illuminating the profound impact of Zn^2+^ binding on caspase-3’s energy landscape. Notably, the Zn^2+^-bound state exhibited a more constrained conformational ensemble, suggesting a reduction in protein flexibility upon Zn^2+^ binding.

**Figure 3.**
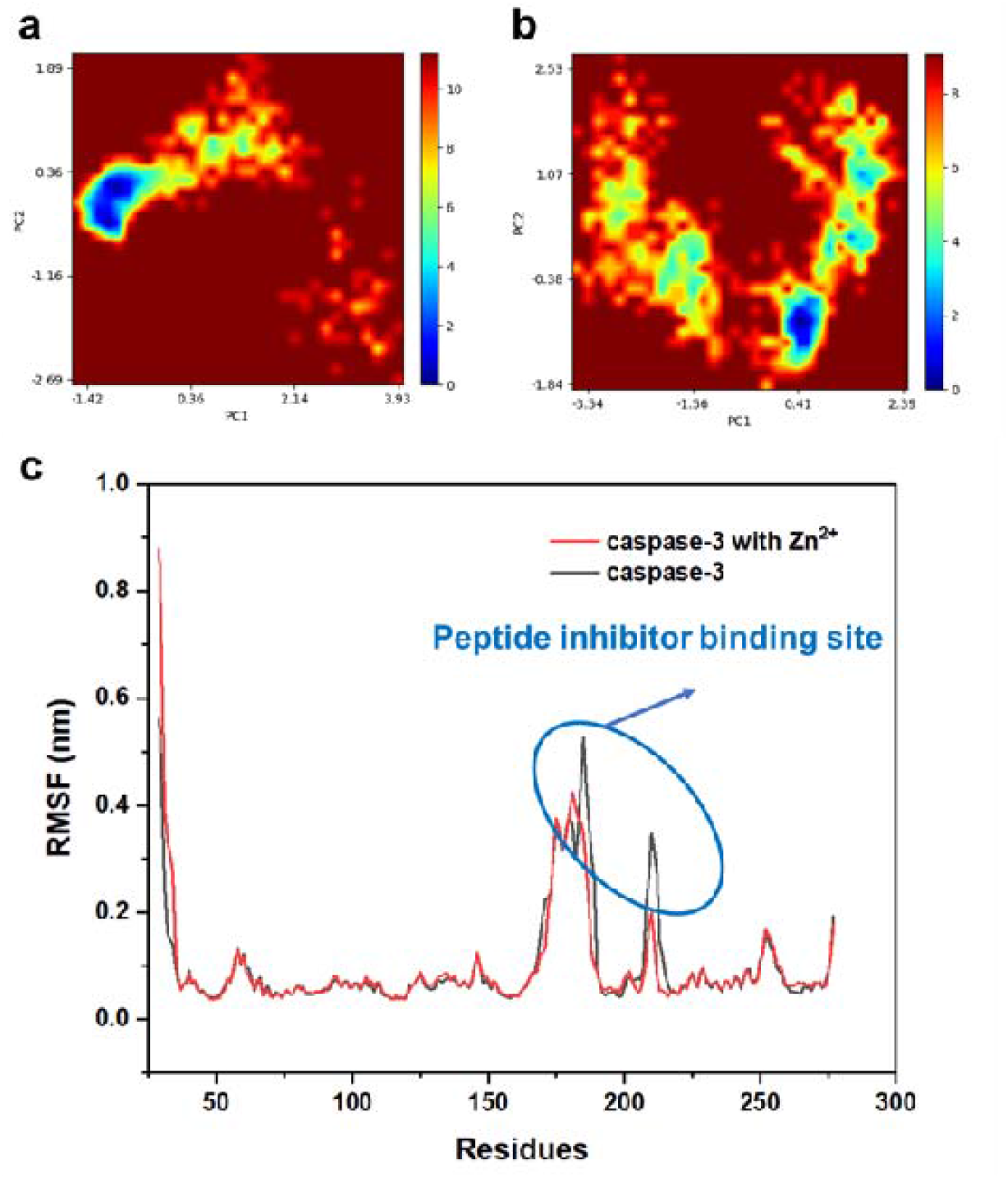
Zn^2+^-mediated modulation of caspase-3 dynamics. (a) Free energy landscape distributions of caspase-3 in the presence (a) and absence (b) of Zn^2+^ ions, revealing distinct conformational preferences. (c) Root-mean-square fluctuations (RMSF) of caspase-3 residues in Zn^2+^-bound (red) and Zn^2+^-free (black) states, highlighting regions of altered flexibility. Key residues involved in Zn^2+^ coordination and catalytic activity are indicated.

To complement this analysis, we conducted a detailed examination of the root-mean-square fluctuations (RMSF) of protein residues across both systems (**Figure 3c**). This comparative RMSF analysis revealed localized changes in structural flexibility, particularly in the vicinity of the active site. Specifically, regions critical for substrate recognition and catalysis showed significantly reduced mobility in the Zn^2+^-bound state. These findings further substantiate the Zn^2+^-induced constraint on peptide binding pocket mobility and adversely influence inhibitory effect of these competitive peptide inhibitors, providing atomic-level insights into the mechanism of Zn^2+^-mediated caspase-3 inhibition. Moreover, we observed non-competitive effects extending beyond the immediate vicinity of the Zn^2+^-binding site, suggesting a complex network of structural perturbations induced by Zn^2+^ binding. These long-range effects may play a crucial role in modulating caspase-3 activity and offer potential targets for non-competitive inhibitor design.

## 3. Conclusion

Our study provides comprehensive insights into the inhibitory mechanisms of caspase-3, a key enzyme in apoptotic pathways. By combining advanced computational techniques, including molecular docking, dynamics simulations, binding free energy calculations, and AlphaFold3 predictions, we have elucidated both competitive peptide inhibitory and Zn^2+^-mediated non-competitive inhibitions to caspase-3. This dual approach offers a more complete picture of caspase-3 regulation. Our analysis of competitive inhibitors demonstrated their effective occupation of the substrate-binding site, providing a structural basis for their inhibitory action. We then identified potential Zn^2+^-binding sites and characterized Zn^2+^-induced conformational dynamics, revealing a mechanism for non-competitive inhibition and its interplay with the competitive inhibition. These findings enhance our understanding of caspase-3 regulation and provide a foundation for rational inhibitor design. The complementary nature of these interplayed inhibition mechanisms suggests potential for synergistic therapeutic approaches. Future studies should focus on experimental validation of these computational insights and explore their

translational potential in developing targeted therapies for apoptosis-related disorders, potentially combining competitive inhibitors with modulators of Zn^2+^-mediated regulation.

