## Supplemental Information for "Computational Analysis of Zn^2+^ Mediated Non-Competitive Inhibition to Caspase-3"

**Table of Content**

**Experimental section**

**Figure S1.** Zinc ion binding sites predicted by alphafold3

**Figure S2.** RMSD values (nm) of the peptide backbone atoms.

**Experimental section**

**Molecular simulation**

The binding sites of zinc ions in caspase-3 were predicted using Alphafold3 sever^1^ with default parameters. Initial peptide structures were generated using the Tleap module of AmberTools^2^, applying standard amino acid parameters. Non-canonical residues were incorporated into the peptides using the open-source version of PyMOL (version 2.5.0), with manual adjustments to sidechain and backbone positions to mitigate steric clashes. Autodock Vina^3^ version 1.2.3) was used to dock the inhibitor peptide into the pocket where DEVD is located in the cocrystal structure of DEVD and caspase-3 protein (PDB ID: 3PCX), using a grid box of 20 × 20 × 20 Å centered on the DEVD binding site. The Amber14SB force field^4^ was employed, with general AMBER force field (GAFF)^5^ parameters assigned for synthetic residues. Partial atomic charges were obtained through RESP fitting in Gaussian 16 (Revision C.01) at the HF/6-31G* level of theory and suitably amended using custom bond, angle and dihedral terms for non-peptide linkages. The Amber ff14SB force field was employed for all molecular dynamics (MD) simulations. The system was neutralized with counterions (Na+, Cl-) and solvated in a TIP3P^6^ water box, maintaining a minimum buffer distance of 10 Å from the protein surface. Prior to the production run, the system underwent energy minimization using a hybrid approach of steepest descent (1,000 steps) and conjugate gradient algorithms (4,000 steps) for a total of 5,000 cycles^7, 8^. Subsequently, a gradual temperature annealing from 10 to 310 K was performed over 0.05 ns under weak harmonic restraints (15 kcal/mol/Å2) on protein-heavy atoms. Density and pressure equilibration were conducted for 1 ns under isothermal-isobaric conditions (310 K, 1 atm) utilizing Langevin dynamics^9^ and Berendsen/Parrinello-Rahman barostats^10, 11^.

Upon confirmation of adequate equilibration (RMSD < 2 Å for protein backbone atoms), 500 ns production runs were initiated using GROMACS 2022.4^12^. with a time step of 2 fs. The LINCS algorithm was used to constrain all bonds involving hydrogen atoms. Long-range electrostatic interactions were treated using the Particle Mesh Ewald method with a cutoff of 10 Å. Molecular dynamics simulations revealed stable binding, with root-mean-square deviation (RMSD) stabilizing within 6 Å after 200 ns.

Subsequent binding energy calculations -- on this trajectory were performed using 400 frames extracted at 500 ps intervals from the last 200 ns of the simulation via the MM/PBSA method. All MM/PBSA calculations were performed by using a modified shell script gmx_mmpbsa^13^. The binding free energy was calculated using the following formula:

$$\Delta G_{binding}=\Delta G_{complex}-(\Delta G_{protein}+\Delta G_{ligand})$$

The ∆G_complex_ represents the protein-ligand complex's overall free energy. The total free energy of the separated protein and ligand in a solvent are represented simultaneously by the ∆G_protein_ and ∆G_ligand_, respectively.

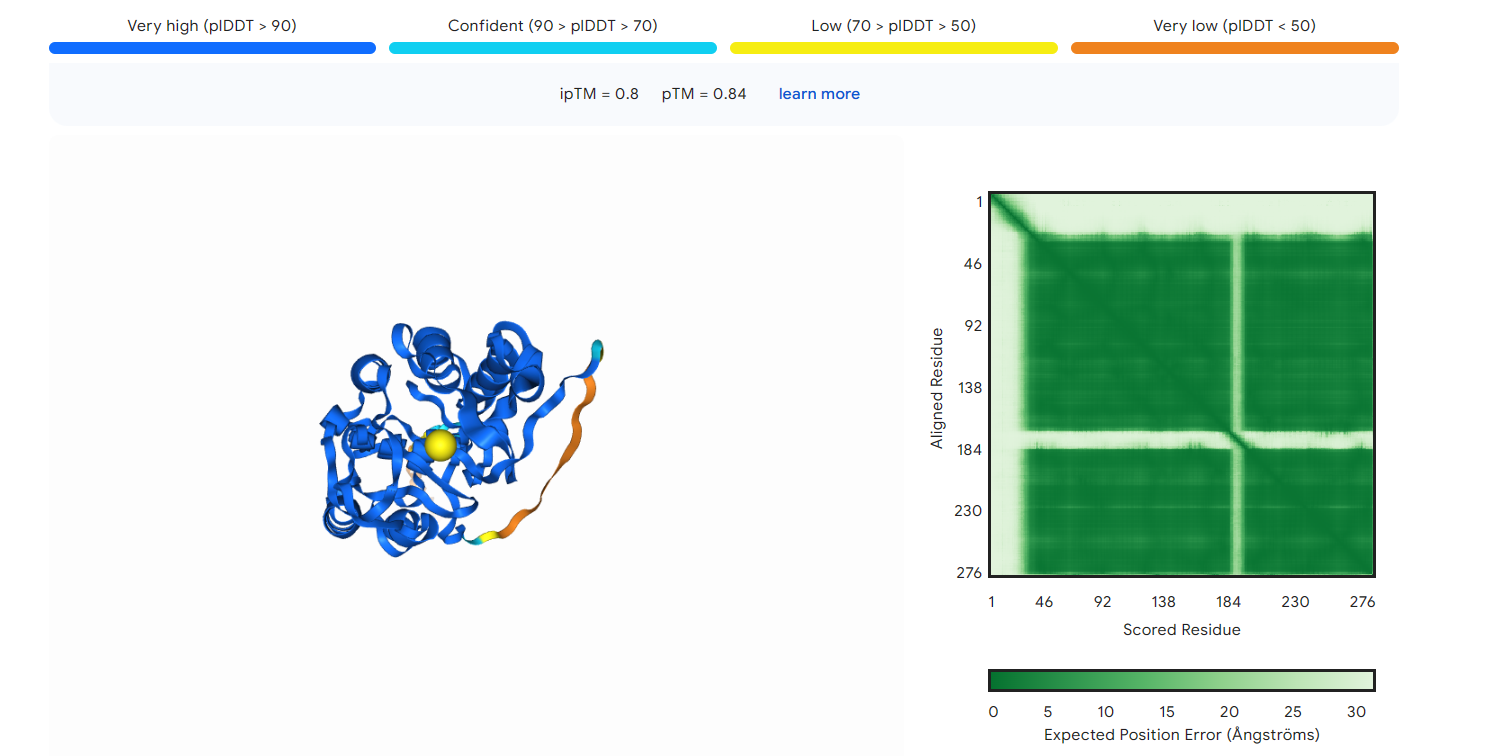

**Figure S1.** Zinc ion binding sites predicted by alphafold3.

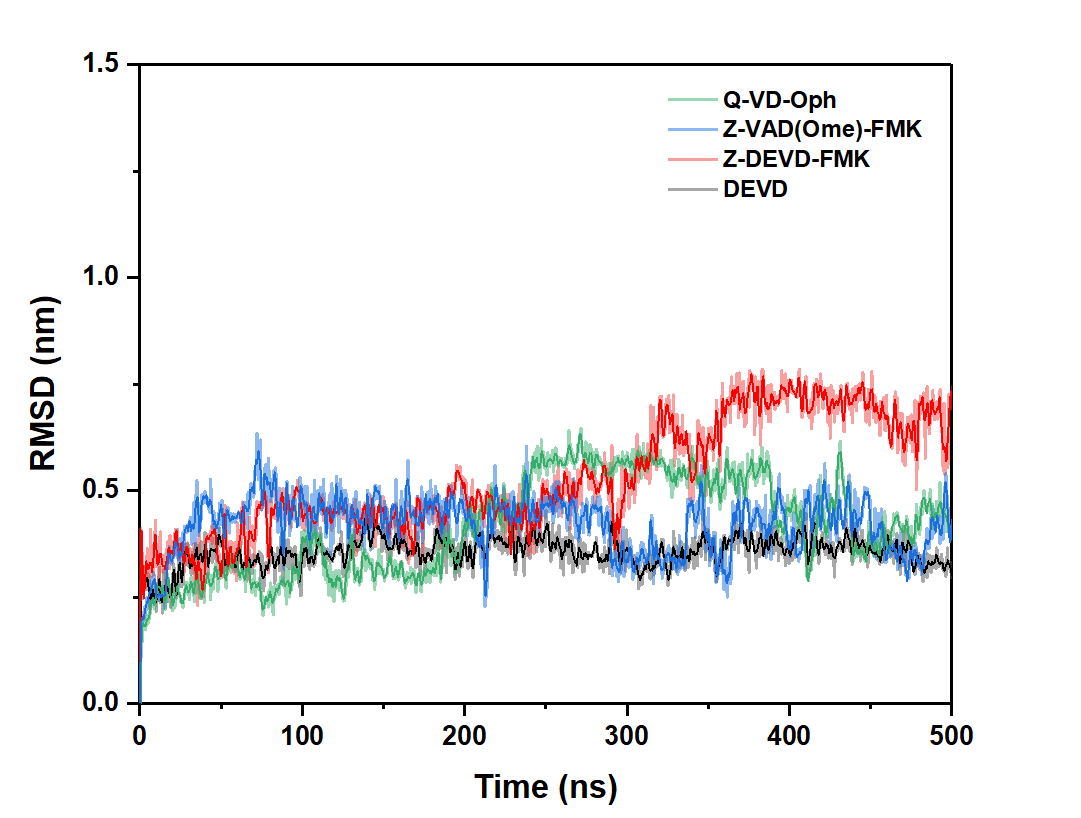

**Figure S2.** RMSD values of the peptide backbone atoms during the 500 ns MD simulations.
